# Acid-Sensing Ion Channels Mediate Type III Adenylyl Cyclase-Independent Acid-Sensing of Mouse Olfactory Sensory Neurons

**DOI:** 10.1101/765420

**Authors:** Juan Yang, Liyan Qiu, Matthew Strobel, Amanda Kabel, Xiangming Zha, Xuanmao Chen

**Author notes:** Correspondence should be addressed to Dr. Xuanmao Chen: Department of Molecular, Cellular and Biomedical Sciences, University of New Hampshire, 389 Rudman Hall, 46 College Road, Durham, NH 03824. These authors contributed equally to the study (Juan Yang, Liyan Qiu).

## Abstract

Acids can disturb the ecosystem of wild animals through altering their olfaction and olfaction-related survival behaviors. It is known that the main olfactory epithelia (MOE) of mammals rely on odorant receptors and type III adenylyl cyclase (AC3) to detect general odorants. However, it is unknown how the olfactory system sense protons or acidic odorants. Here we show that the mouse MOE responded to acidic volatile stimuli in the presence and the absence of AC3. Acetic acid-induced electro-olfactogram (EOG) responses in wild type (WT) MOE can be dissected into two components: one dependent on the AC3-mediated cAMP pathway and the other not. MOE of AC3 knockout (KO) mice retained an acid-evoked EOG response but failed to respond to an odor mix. The acid-evoked responses of the AC3 KO could be blocked by diminazene, an inhibitor of acid-sensing ion channels (ASICs), but not by forskolin/IBMX, which desensitize the cAMP pathway. AC3 KO mice lost their sensitivity to detect pungent odorants but maintained sniffing behavior to acetic acid. Immunofluorescence staining demonstrated that ASIC1 proteins were highly expressed in olfactory sensory neurons (OSNs), mostly enriched in the knobs, dendrites, and somata, but not in olfactory cilia. Moreover, mice exhibited reduced preference to attractive objects when placed in an environment with acidic volatiles. Together, we conclude that the mouse olfactory system has a non-conventional, ASICs-mediated mechanism for acid-sensing. Acid stimulation of ASICs may unselectively depolarize different OSNs and interfere with the anatomical logic for odor perception.

## Introduction

Protons in volatile or dissolved acids are the simplest odorants that an animals’ olfactory system typically encounter in the environment. Acids affect ecological stability and influence animals’ key survival behaviors including food discrimination, avoiding predator, and host-seeking [1-3]. Salmon can lose the ability to smell dangerous predators as oceans become more acidic due to increasing carbon emission [4]. The ability for sea bass’s to sense and respond to odors of predators and food sources is more strongly influenced by acidified water than by other odors [1]. Fruit flies sense the concentrations of acetic acid to discriminate the ripeness of food [5]. Additionally, acidic volatiles in human sweat stimulate mosquitoes’ olfactory system and help them identify a host for blood-feeding [2]. Furthermore, human subjects who are repetitively exposed to acetic acid in their home environment exhibit decreased sensitivity to chemical irritancy [6]. Despite the prevalence of acidic volatiles in the environment and their ecological impact, the molecular mechanisms of acid-sensing in the vertebrate olfactory system is still poorly understood.

Acidic volatiles, such as acetic acid, inflowing into the nasal cavity are dissolved in the humidified nasal mucosa and then bind to their receptors and stimulate OSNs [7, 8]. Depending on the acids’ strength, some acidic volatiles such as acetic acid can dissociate into protons (H+) and bases (acetate) in the nasal mucosa. It is known in mice, that OSNs rely on olfactory receptors and AC3 to detect regular odorants [9-12]. Olfactory receptors in olfactory cilia are instigated by odorants, stimulating the olfactory G-protein subunit (G_olf_), which subsequently activates AC3 to generate cAMP. This in turn opens cyclic nucleotide gated (CNG) channels, leading to cation influx and depolarization of OSNs [13-16]. The un-dissociated acid and the base from acidic volatiles most likely bind to regular olfactory receptors in olfactory cilia. Nevertheless, it is unknown how protons in the nasal mucosa affect OSNs. In this study, we aimed to determine which receptors in OSNs the protons bind to, whether the AC3-mediated cAMP pathway is required for acid-sensing, and in a broader scope whether acidic volatiles affect normal olfactory perception.

Recently, two ionotropic receptors (IR), IR8a and IR64a, have been identified in the insect olfactory system [2, 17], and they directly mediate the acid-sensing of mosquitoes and Drosophila. Interestingly, all known olfactory receptors of insects are ionotropic [18], in sharp contrast to the metabotropic olfactory receptors of mammals. We initially observed that the MOE of AC3 KO mice retained the sensitivity to acidic volatiles in EOG recordings (see Figure 1). This led us to hypothesize that acid-sensing by the mammalian olfactory system may be independent of the olfactory receptor- and AC3-mediated metabotropic pathway, but instead utilize the primitive ionotropic mechanism, as insects do. We further reasoned that ASICs are ideal candidates for the acid-sensing of OSNs. This is because ASIC1a-containing ASICs are the most pH-sensitive ion channels that have been identified [19, 20], and ASIC currents have been detected in virtually every neuron in the brain [21]. ASICs contain a high abundance of charged amino acids (glutamate and aspartate), which are exquisitely assembled in their ectodomain [22]. This confers ASICs with a high sensitivity to detect subtle pH variation. More relevantly, ASIC1 mRNA has been detected in OSNs using an RNA-Seq approach [23] and a quantitative real time (qRT)-PCR method [24]. In addition, ASIC1 has recently been reported to regulate normal olfactory function [25] but its mechanism remains unknown. Here we show that mouse MOE can respond to acid air puffs in the presence and the absence of AC3. Acidic volatiles-induced EOG responses in AC3 KO samples were completely blocked by diminazene, an ASIC inhibitor [22, 26]. ASIC1 was highly expressed in OSNs in the mouse MOE; it was highly distributed in the knobs, the dendrites and the somata, but not in olfactory cilia. In addition, wild type mice had a reduced preference to attractive objects when placed in an acidic volatile environment. These results indicate that the acid-sensing by mouse MOE is independent of the metabotropic receptor and AC3-mediated pathway. Instead, it is directly mediated by ASICs, ionotropic proton-gated receptors. Given ASICs are much more widely distributed in the MOE than individual types of olfactory receptors, ASIC activation may unselectively cause depolarization of OSNs and interfere with the anatomical logic of odor perception.

**Figure 1.**
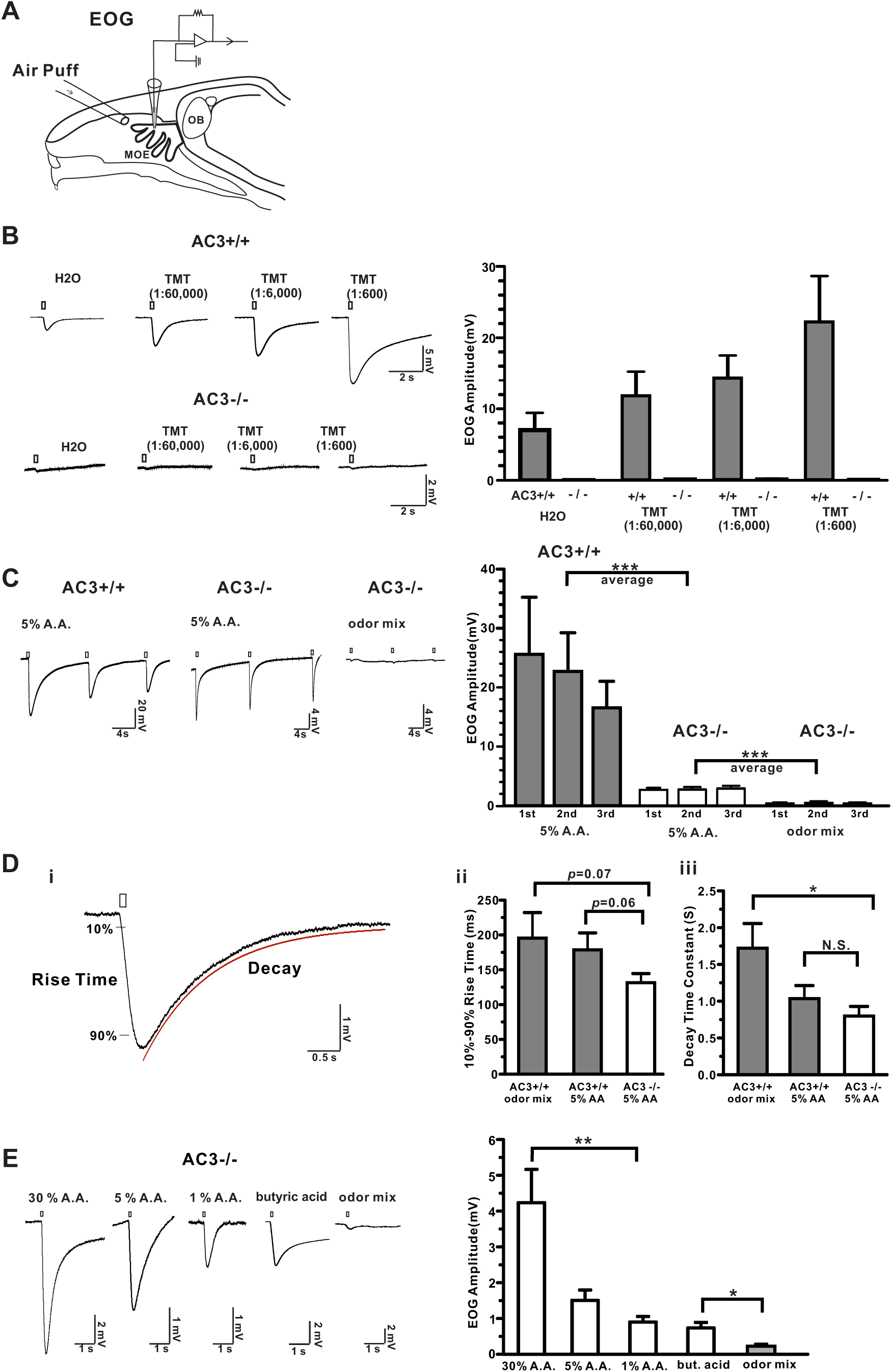
The MOE of AC3 KO mice lost the sensitivity to regular odors but maintained responses to volatile acids. (A) A diagram of EOG recording. (B) The MOE of AC3 KO mice lost the sensitivity to pungent TMT in EOG recording. Left, representative traces of TMT-elicited EOG responses in MOE of AC3 WT (n = 6) and AC3 KO mice (n = 6). Right, statistical summary of TMT-evoked EOG responses. (C-E) MOE of AC3 KO mice retained a sensitivity to volatile acids. (C) Acetic acid (A. A., 5%) elicited pronounced EOG responses in MOE of both AC3 WT and KO mice. A odor mix failed to elicit EOG responses in MOE of AC3 KO mice. Left, representative traces; Right, statistical data of EOG amplitude, n = 3-6. *** *p* < 0.001, comparing averages of 3 repeated EOG measurements with unpaired Student’s t-test. (D) The 10%-90% rise time and decay time constants of acetic acid- and odor-induced EOG responses. (i) A representative trace showing how 10%-90% rise time and decay time constants were measured. n = 6-11. (ii) Statistical bar graph of 10%-90% rise time. An ANOVA test yielded no significance among the three groups. However, low *p* values were generated from unpaired Student t-test between two groups. (iii) Statistical bar graph of decay time constants. ANOVA with post hoc Turkey’s multiple comparison test, F (2, 22) = 5.2, * *p* < 0.05. (E) AC3 KOs’ MOE retained the olfactory sensitivity to various concentration of acetic acids as well as to volatile butyric acid (concentration, 2 M). Left, acid-elicited EOG responses in MOE of AC3 KO mice. Right, statistical bar graph of EOG amplitude. n = 6-9. ** *p* < 0.01 with ANOVA test, F (2, 18) = 6, comparison of 3 acetic acid concentrations. * *p* < 0.05, butyric acid Vs. odor mix, using unpaired student’s t-test.

## Results

### The MOE of AC3 KO mice lose sensitivity to general odorants but retain responses to acidic volatiles

AC3 represents an essential enzyme in mediating the main olfactory signal transduction pathway of mammals, and ablation of AC3 leads to anosmia - loss of smell [12]. We first used AC3 WT and KO MOE samples in EOG recording to examine their olfactory sensitivity to regular odorants and acidic volatiles. To confirm that AC3 KO mice lost olfactory sensitivity, even to pungent odorants, we recorded EOG in response to 2,4,5-trimethylthiazoline (TMT), a pungent predator odor of fox feces [22, 27], using AC3 WT and KO MOE samples. WT MOE samples had pronounced EOG responses to different concentrations of TMT. However, AC3 KO samples failed to yield any responses to TMT (Fig. 1B), confirming that AC3 KO mice lost olfactory sensitivity even to a pungent odorant. Next, we puffed 5% acetic acid and non-acidic odor mix to MOE samples of AC3 WTs and KOs, respectively. Surprisingly, we found that acetic acid could elicit pronounced EOG responses in both AC3 WT and KO samples, although the EOG amplitude recorded in KO MOE were much smaller than that of WT. Consistently, an odor mix failed to evoke any responses in KO samples (Fig. 1C). The kinetics of acetic acid-evoked EOG traces was different in WT and KO samples. The acetic-acid evoked EOG amplitude in WT samples exhibited certain rundown over repetitive stimulation, while responses in KO samples did not (Fig. 1C). Both the 10%-90% rise time and the decay time constants of the acetic acid-evoked EOG responses in KO MOEs were slightly smaller than those of odor mix-evoked and acetic acid-evoked EOG responses of WT MOEs (Fig. 1D). Additionally, to confirm that the responses were caused by acidic volatiles and not by mechanical artifacts, we puffed different concentrations of acetic acid and butyric acid (another volatile acid) to test the EOG responses in AC3 KO samples. Acetic acid-elicited EOG responses were dependent on acetic acid concentrations. Air puffs of butyric acid also evoked a good EOG response, which was much higher than the odor mix-evoked responses (Fig. 1E). These data indicate that the mouse MOE has an AC3-independent mechanism to sense acidic volatiles.

### ASICs mediate AC3-independent acetic acid-induced EOG responses

To validate that acid-elicited EOG responses are not mediated by the AC3-mediated cAMP pathway, we used forskolin/IBMX, which can activate and subsequently desensitize the olfactory cAMP pathway [28]. We found that both acetic acid and odor mix could elicit strong EOG responses in WT samples (Fig. 2A). After treating with forskolin/IBMX, the odor mix-evoked EOG responses were abolished, whereas the acetic acid-evoked responses persisted with a markedly reduced amplitude (Fig. 2A). Moreover, the acetic acid-evoked EOG responses recorded in AC3 KO MOE were not affected by forskolin/IBMX (Fig. 2B). Kinetically, the rise time and decay time constants of acetic acid-evoked EOG responses recorded from AC3 KO samples were comparable to those of EOG responses recorded in AC3 WT samples that had been pre-treated with forskolin/IBMX (Fig. 2C). These results corroborate the concept that the acid-evoked EOG response is mediated by some mechanism independent of the AC3-mediated cAMP pathway. ASICs are known to be the most pH-sensitive ion channels and are widely distributed throughout the nervous system [19, 21, 22, 26, 28-31]. ASIC1 mRNA is detected in the mouse MOE by qRT-PCR experiments [24] and ASIC1a’s mRNA is found in OSNs using a single OSN RNA sequencing technique [23]. In contrast, the transcript of transient receptor potential channel V1 (TRPV1), a less pH-sensitive ion channel, has not been detected in OSNs in the same RNA-Seq experiments [23]. Hence, we postulated that ASICs mediate the AC3-independent component of acid-sensing by mouse OSNs. To test this, we used diminazene, a potent ASIC blocker [26], in the EOG recording using AC3 KO MOE samples. Indeed, application of diminazene abolished the acetic acid-evoked EOG responses (Fig. 2D), which was partially reversible after washing away. These results indicate that the acid-evoked EOG response is directly mediated ASICs, independent of the AC3-mediated cAMP pathway.

**Figure 2.**
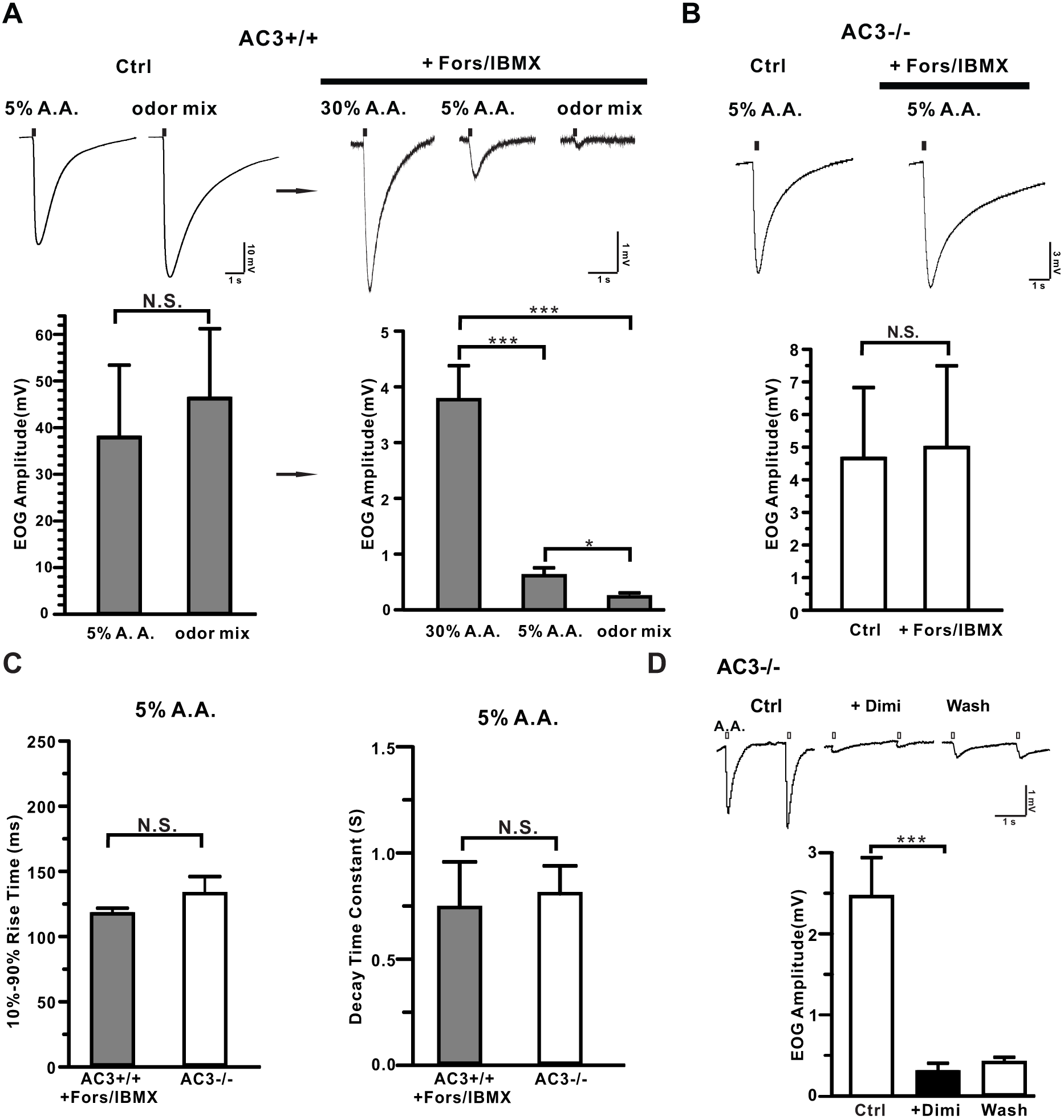
Acetic acid-induced EOG responses were dissected into two components: AC3-dependent and AC3-independent components. (A) Forskolin/IBMX eliminated odor-induced EOG responses, but not acetic acid-induced responses in WT mice. Top, representative traces of EOG recording on MOE of AC3 WT mice. Left, control EOG recording. Both 5% acetic acid and odor mix stimulated high EOG responses. Right, application of forskolin/IBMX eliminated odor responses, but not acetic acid-response. Bottom, statistical bar graphs of EOG amplitude under different conditions. Note the change of EOG amplitude scale from the left graph to the right. Left, n. s., not significant by unpaired Student’s t-test. Right, acetic acid still evoked pronounced EOG responses after foskolin/IBMX treatment. ANOVA with Post-hoc Turkey’s multiple comparison, F (2, 16) = 25, *p* < 0.0001, n = 6-7. (B) Acetic acid-evoked EOG responses in AC3 KO MOE samples were insensitive to forskolin/IBMX treatment. Top, representative EOG traces evoked by 5% acetic acid. Bottom, statistical bar graph of EOG recording. n = 3, n. s., not significant with unpaired Student’s t-test. (C) Statistical bar graph of 10%-90% rise time and decay time constant of acetic acid-evoked EOG responses in AC3 WT MOE after forskolin/IBMX treatment (n=6) compared with those of AC3 KOs (n = 11), n. s., not significant by unpaired Student’s t-test. (D) Acetic acid-evoked EOG responses in the MOE of AC3 KOs were blocked by diminazene (Dimi, 200 µM). Left, representative traces of EOG recording. Right, statistical bar graph of EOG recording, n= 5, *** p< 0.001.

### AC3 KO mice lose sensitivity for regular odorants but retain the ability to detect acidic volatiles behaviorally

Next, we conducted a three-chamber avoidance test and a Q-tip cotton swab habituation/dishabituation test to determine whether AC3 KO mice lost the sensitivity to detect a pungent odorant but still retained the ability to sense acidic volatiles behaviorally. In the 3-chamber avoidance test, we placed a TMT-moisturized object in the right chamber and a control object in the left, and allowed subject mice to freely explore the three chambers. AC3 WT mice spent much less time in the chamber with TMT than the other two chambers (Fig. 3A), suggesting that AC3 WT mice could detect the predator odor and attempted to avoid it. In contrast, the times that AC3 KO mice stayed in each chamber did not show significant differences (Fig. 3A), indicative of the loss of sensitivity to TMT. In a Q-tip habituation/dishabituation test, we placed a Q-tip which carried water vehicle into the mouse’s home cage repetitively 4 times, followed by a Q-tip carrying TMT odor the fifth time and a vehicle Q-tip the sixth time. The sniff time at the fifth repetition increased significantly for AC3 WT mice, but not for AC3 KO mice (Fig. 3B). Interestingly, when we used a Q-tip moisturized by 5% acetic acid at the fifth time, both AC3 WT and KO mice exhibited a strong interest to sniff the Q-tip (Fig. 3C). These data demonstrate that AC3 KO mice lose the olfactory sensitivity to pungent odors but retain the ability to detect volatile acid behaviorally.

**Figure 3.**
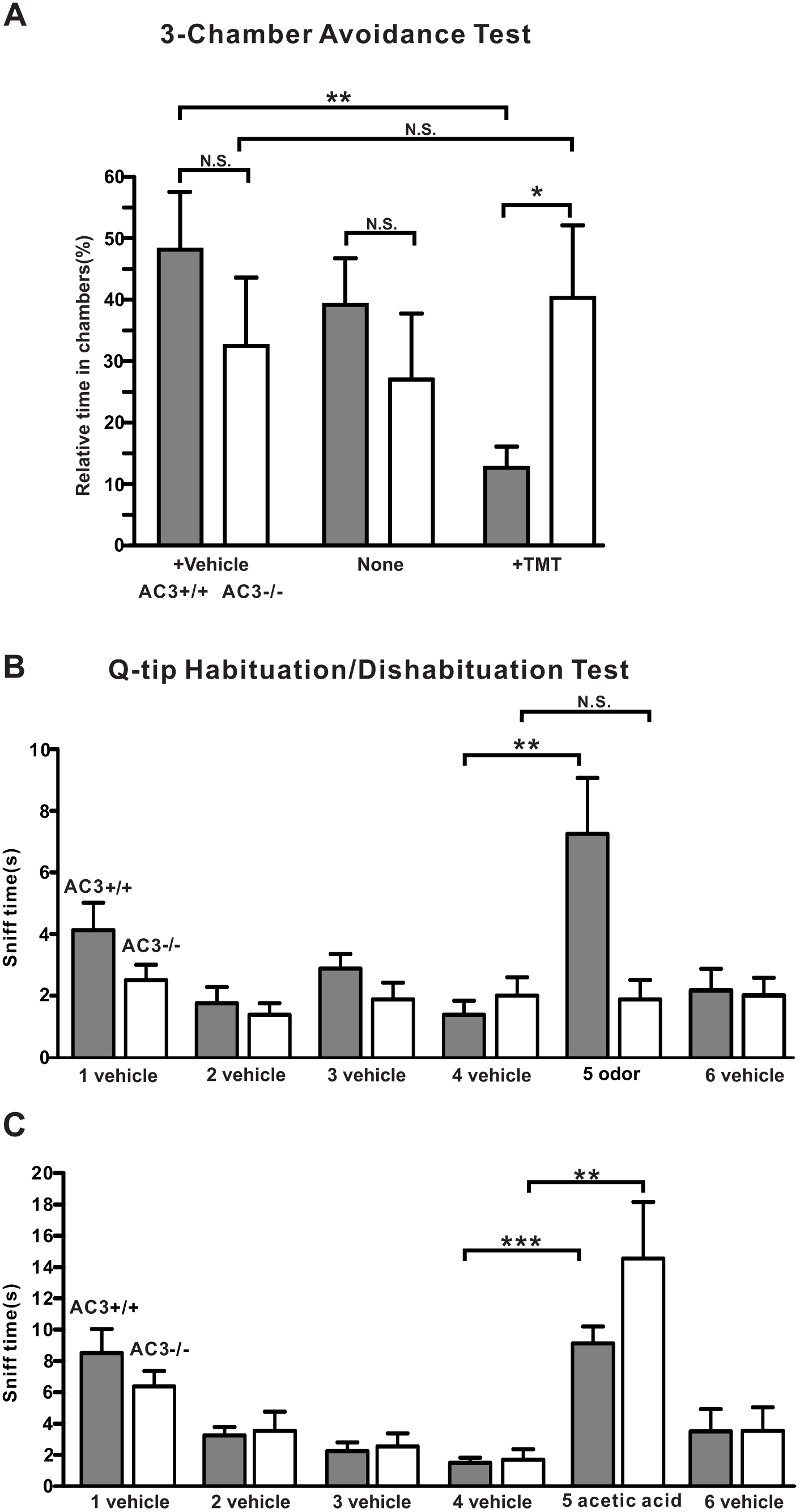
AC3 KO mice lost olfactory sensitivity to pungent odorants, but still exhibited sniffing responses to acetic acid volatiles. (A) 3-chamber TMT avoidance test using AC3 WT and AC3 KO mice. Top, configuration of 3-chamber avoidance test. The left and right chambers were placed with TMT and vehicle respectively. Bottom, statistical bar-graph of relative time in each chamber. AC3 WT mice spent more time in the chamber with vehicle and much less time in the chamber containing TMT. AC3 KO mice showed no preference to any chambers. n = 7 pairs of WTs and KOs. (B) Q-tip habituation/dishabituation test at home cage. The sniff time to TMT Q-tip (the fifth Q-tip) increased significantly in AC3 WT mice, but no significant change in AC3 KO mice in the test. Data were collected from 8 WTs and 8 KOs, ** *p* < 0.01 n. s. not significant by paired Student’s t-test. (C) In the habituation and dishabituation test, TMT Q-tip was replaced by 5% acetic acid Q-tip. Both AC3 WT and KO mice increased sniff time significantly at the fifth time (the acetic acid Q-tip). AC3 KO adult mice retain sniffing responses to acids. Data collected from 8 WTs and 6 KOs. ** *p* < 0.01, ***, *p* < 0.001, with paired Student’s t test

### ASIC1 protein is expressed in the somata, dendrites, and knobs of OSN, but not in olfactory cilia

It is unknown whether ASIC proteins are present in the OSNs of mouse MOE and if so, where they are expressed. We used ASIC1 KO MOE samples to validate the specificity of several anti-ASIC antibodies. We successfully identified one monoclonal anti-ASIC1 antibody (#75-277, UC Davis NeuroMab Facility) that yielded a clear immunostaining signal in the MOE of WT mice, but not in ASIC1 KO mice (Fig. 4A), indicative of a high specificity of the antibody against ASIC1. Next, we co-stained this antibody with AC3 antibody in immunostaining to probe the ASIC1 expression pattern using ASIC1 WT and KO, AC3 WT and KO samples. We observed that ASIC1 was predominantly expressed in the OSN layer in ASIC1 WT mice throughout the MOE (Fig. 4A&B), but not in the MOE of ASIC1 KOs (Fig. 4A). Interestingly, ASIC1 immunostaining signals were mostly detected in the knobs, the dendrites, and somata of OSNs, but did not overlap with AC3 in olfactory cilia (Fig. 4C-ii). Some ASIC1-labeled knobs localized in the proximity of the mucus layer, even if they were not overlapped by AC3 signals (Fig. 4C-i). This expression pattern allows ASIC1-expressing knobs to be accessible by protons dissociated from volatile acids. Additionally, ASIC1 was highly expressed in the tip of WT’s MOE turbinates, where AC3’s expression was absent (Fig. 4D top), and abundantly enriched in some processes facing toward the mucosal cavity. This ASIC1 staining pattern was absent in ASIC1 KO samples (Fig. 4D bottom). This result further supports the possibility that ASIC-mediated acid-sensing is independent of AC3. We estimated the proportion of ASIC1-positive OSNs in the MOE. ASIC1 expression varied a bit throughout the MOE and the average percentage was about 6.6±1.1 % (Fig. 4E). ASIC1 immunostaining signals were also detected in a few sustentacular cells, but at a much lower percentage (Fig. 4C-iii). The ratio of ASIC1-positive signal in sustentacular cells relative to ASIC1-positive signals in OSNs was estimated to be 0.03±0.01 (data collected 9 MOE areas from 3 animals). The expression abundance of ASIC1 proteins in AC3 KOs was lower than in WTs (Fig. 4F). Collectively, the expression pattern of ASIC1 in OSNs allows ASIC1 to directly mediate the acid-sensing in the MOE.

**Figure 4.**
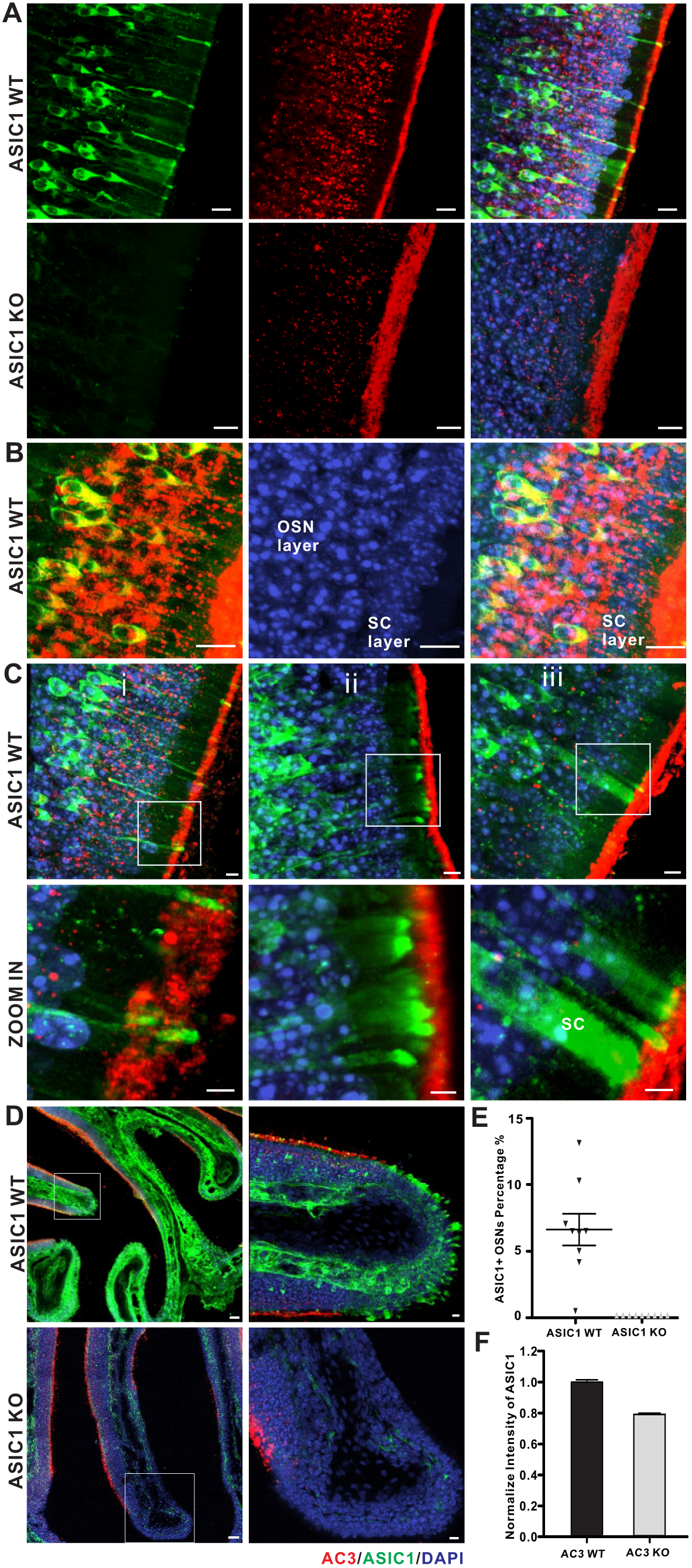
ASIC1 expression pattern in the mouse MOE. (A) Immunostaining using antibodies against ASIC1 (green), AC3 (red), together with DAPI (blue) on MOE samples from ASIC1 WT (top) and KO mice (bottom). ASIC1 immunostaining signals were detected in the MOE of ASIC1 WT mice, not ASIC1 KO mice. Scale bar, 10 µm. (B) Double immunostaining with antibodies against AC3 (Red), ASIC1 (green) and DAPI (blue) on ASIC1 WT MOE samples. ASIC1 staining signals were mostly found in the OSN layer and overlapped with AC3 in the cell body of OSN. AC3 staining signals were boosted in order to differentiate OSN layer with supporting cell layer. Scale bar, 10 µm. (C) (i-ii) ASIC1 proteins were enriched in the knobs (yielded the strongest staining), dendrites, and somata of OSNs, but not overlapped with AC3 staining in olfactory cilia. (iii), ASIC1 proteins were also detected in a few sustentacular cells (SC). Boxes in pictures on the top were zoomed in and shown on the bottom. Scale bars: top 5 µm; bottom 3 µm. (D) Special ASIC1 expression pattern at the tip of turbinates. Top, ASIC1 proteins were abundantly expressed at the tip of tubinates in WT mice where AC3 was absent. Bottom, similar structures were found, but no ASIC1 staining signal detected in ASIC1 KO samples. Boxes in pictures on the left were zoomed in to pictures on the right. Scale bar, 100 µm (left), 10 µm (right). (E) Estimation of ASIC1-positive neurons among OSNs in ASIC1 WTs’ and KOs’ MOE samples. A statistical bar graph shows the distribution percentage of ASIC1-positive neurons normalized to total OSN number. Data was measured in 9 randomly chosen areas out of 3 mice’s MOE. The average distribution percentage of ASIC1 positive OSNs is 6.6 ± 1.1 % in ASIC1 WT mice, and 0% in ASIC1 KO mice. (F) ASIC1 intensity in the mouse MOE of AC3 WT and KO mice. The expression of ASIC1 proteins was decreased in the MOE of AC3 KO mice. WTs’ and KOs’ imaging signals were normalized to their own DAPI staining. Images from at least 9 regions out of 3 mice were quantified.

Moreover, homotrimeric ASIC1 and heterotrimeric assemblies of distinct ASIC subtypes (ASIC1-3) are both functional channels in neurons [32-34]. To examine whether other ASIC subtypes are expressed in the mouse MOE, we tested an anti-ASIC2 antibody (Catalog# ASC-012, Alomone Labs). However, this anti-ASIC2 antibody yielded unspecific signals in both ASIC2 KO and WT samples, so we failed to verify the expression of ASIC2 protein in OSNs. Nevertheless, ASIC2 mRNA has been detected in a qRT-PCR study [24]. In addition, we also used ASIC3 antibodies (Catalog#: ASC-018, Alomone Labs) to stain the mouse MOE. However, our staining signal was negative, thus we also failed to verify whether ASIC3 is expressed in OSNs, even if ASIC3 is a sensory neuron-specific subtype [35].

### Acidic volatiles interfere with mouse olfaction

To test whether volatile acids affect mouse olfaction, both WT male and female mice were subjected to a three-chamber acid-interference olfactory test (Fig. 5A). Two different odor objects were placed inside the apparatus, one in the left chamber and the other in the right. For the test using male mouse subjects, we used cotton nestlet that came from adult female mouse cages as an attractive object, and a clean cotton nestlet of the same size as a control. For the test using female mice, we used peanut butter flavoring on filter paper as an attractive odorant object and water filter paper as a control. We compared the effect of different environments on odor-sniffing preference. In an environment without acidic volatiles, male mice spent more time in sniffing female cotton nestlet than clean cotton nestlet. The addition of water (Fig. 5B) or ethyl vanillin (Fig. 5C) to each cotton nestlet did not affect male mice’s preference to female bedding. Similarly, female mice spent more time in sniffing peanut butter than water in a neutral environment (Fig. 5D). However, in an environment with acidic volatiles, the sniffing preferences were affected. When acetic acid was added next to both objects, the preferences for both male mice and female mice spent in sniffing attractive objects were significantly reduced (Fig. 5E-G). These data suggest that acidic volatiles interfere with normal odor detection of mice, whereas ethyl vanillin as a non-acidic odorant does not.

**Figure 5.**
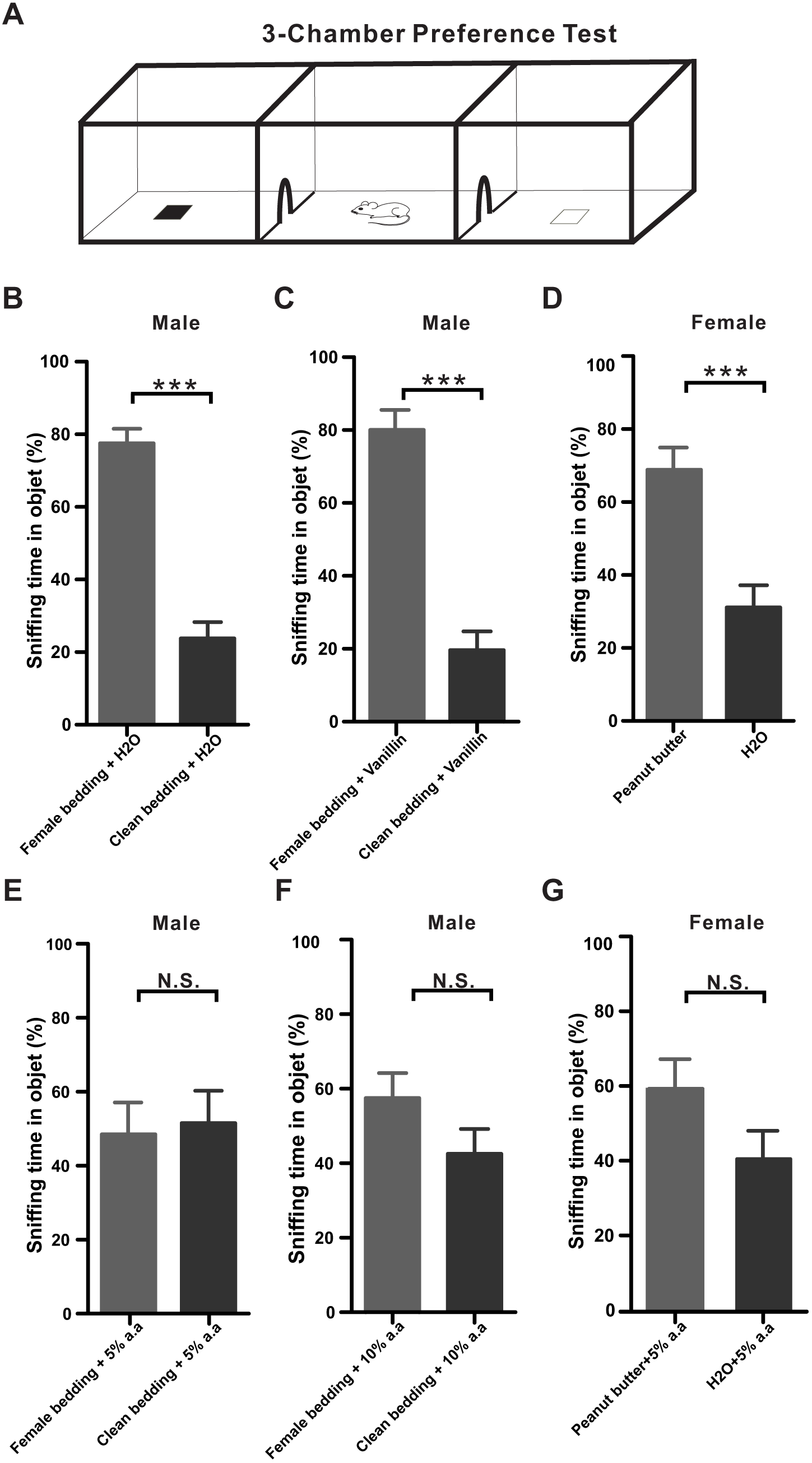
Acidic volatiles interfere with mouse olfaction. (A) A schematic depicting a mouse in 3-chambers behavior test. (B-D) Male mice preferred to sniff female bedding, and female mice preferred to sniff peanut butter in non-acidic volatile environments. (B) Male mice preferred to sniff female cotton nestlet over clean cotton nestlet in the presence of water (20 µl), which moisturized both cotton nestlets. *** *p* < 0.001, n = 8. (C) Male mice still preferred to sniff female cotton nestlet over clean cotton nestlet in the presence of ethyl vanillin (20 µl), which moisturized both beddings. *** *p* < 0.001, n = 8. (D) Female mice preferred to sniff peanut butter flavor over water on filter paper. Each filter paper had additional 20 µl water. *** *p* < 0.001, n = 7. (E-G) The mouse normal olfaction was affected in volatile acid environments. (E) Male mice did not prefer to sniff female cotton nestlet over clean bedding in the presence of 20 µl 5% acetic acid. n.s., not significant, n = 8. (F) Male mice did not prefer to sniff female cotton nestlet over clean bedding in the presence of 20 µl 10% acetic acid. n. s., not significant, n = 8. (G) Female mice did not prefer to sniff peanut butter flavor over water in the presence of 20 µl 5% acetic acid. n. s., not significant, n = 7. All statistic tests were conducted using two-tail unpaired Student’s t-test.

## Discussion

Due to increasing CO2 emission and exacerbated air pollution, humans and wild animals are more frequently exposed to acidic environments. Through affecting the olfactory system, acids disturb the ecosystem and interfere with the survival behaviors of wild animals by impeding their ability to sense predator or food odorants [1-3]. Recent studies have shown that acid-sensing in insects is mediated by IR8a and IR64a, two ionotropic receptors [2, 17]. Yet, it is unknown how acidic volatiles and protons activate OSNs of mammals and influence their olfactory communication. In this study, we sought to understand which molecular mechanism mediates acid-sensing in the mouse olfactory system and examine if acidic volatiles interfere with normal olfaction. This report has presented three major findings: (1) acetic acid-evoked EOG responses in the MOE of WT mice can be dissected into two components: one dependent on the canonical AC3-mediated cAMP pathway and the other not; 2) ASIC1 is highly expressed in the dendrites, the knobs, and the somata but not in the olfactory cilia, and mediates the AC3-independent mechanism for acid-sensing, (3) Acidic volatiles interfere with normal olfaction. By revealing the key molecular mediators in acid sensing in mouse OSNs, our data help to explain how acidic volatiles interfere with normal olfaction in mammals.

The canonical AC3-mediated cAMP pathway mediates the stronger component of the acetic acid-evoked EOG responses. We speculate that this is due to the fact that olfactory cilia harbor all essential signal-transduction proteins including olfactory receptors[9], G_olf_, AC3 [12], and CNG [13] as well as a calcium-activated chloride channel [27], allowing for amplification of the signaling. Consistent with the canonical AC3-mediated cAMP signaling, this strong component was desensitized and abolished by forskolin/IBMX and did not retain in AC3 KO samples (Fig. 2). The AC3-independent EOG component was much weaker than the AC3-mediated component, most likely due to lack of a signal amplification mechanism.

Although mammals mostly utilize metabotropic receptors to detect odorants, the ionotropic pathway for acid sensing may have an ancient origin. Insects almost exclusively utilize ionotropic receptors to mediate olfactory and gustatory senses [18, 36-38]. IR8a and IR64a, which directly mediate the acid-sensing of mosquitos and drosophila, have been reported in the insect olfactory system [2, 17, 39]. In this regard, it is not surprising to find that protons, one of the simplest and antient chemical cues [40] also bind to an ionotropic receptor in mammalian OSNs [30]. Our results strongly suggest the possibility that the second component of acid sensing in the mouse olfactory system is directly mediated by ASICs (Fig. 6), which are proton-gated sodium channels and the most pH-sensitive ion channels [30]. Diminazene, a potent ASIC blocker, completely abolished the acetic acid-evoked EOG responses (Fig. 2D). ASIC1 proteins were found to be highly expressed in WT mouse MOE (Fig. 4A-C), not in ASIC1 KO samples. Intriguingly, ASIC1 is not expressed in the olfactory cilia. Rather, it is enriched in the knobs, dendrites and the somata of OSNs, particularly the knobs (Fig. 4C). The expression pattern of ASIC1 corroborates the concept that the acid sensing does not depend on the AC3-medicated cAMP signaling in olfactory cilia.

**Figure 6.**
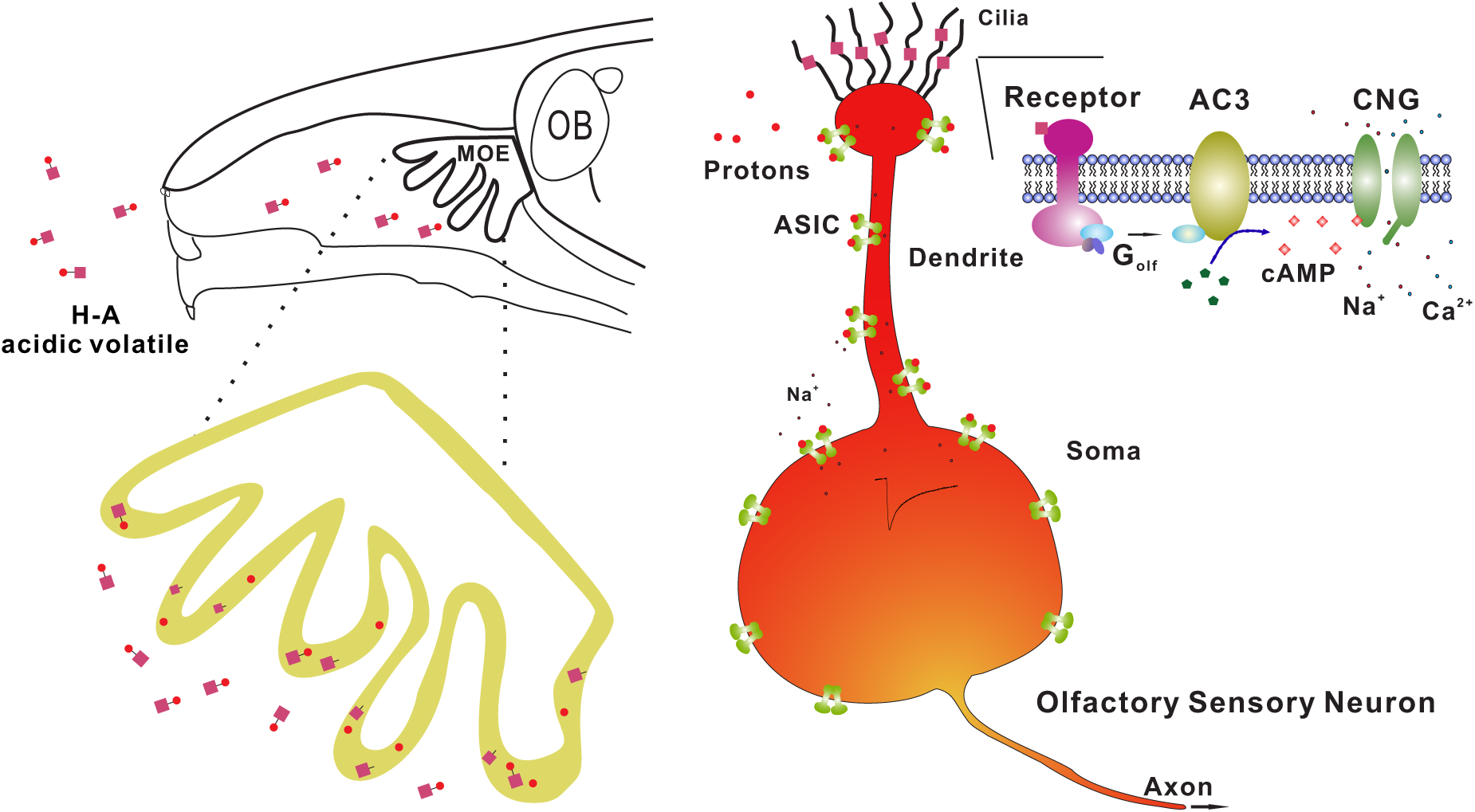
Proposed mechanism of acid-sensing in the mouse MOE. Top, a schematic of acidic volatiles in the air inhaled into the mouse nasal cavity. Bottom, the MOE on the top enlarged to the bottom with single OSN shown to the right. Acidic volatiles (e.g., acetic acid) are dissolved in mucosa and dissociated into protons (H+) and base (acetate). Based on our data, we propose that OSNs in the MOE can be stimulated by protons and base separately. While the base moiety initiates the conventional AC3-medaited cAMP pathway in olfactory cilia, protons directly activate ASICs expressed in the knob, dendrite, and soma, promoting the depolarization of OSNs. As ASICs are more widely expressed than individual subclass of receptor-specific OSNs, ASIC activation may unselectively depolarize different subclasses of OSNs, interfering with the anatomical logic of neural information transmission for specific odorants.

We have estimated that the ASIC1 distribution percentage among OSNs in the MOE is about 6.6%. This number is much higher than the individual olfactory receptor’s expression ratio in the MOE, which generally ranges from 0.1-1.0% [41]. Of note, the ASIC distribution percentage in the MOE could be higher, because we did not count in the contributions of ASIC2, and ASIC3 which is a sensory neurons-specific ASIC subtype [35], due to lack of a specific high-quality antibody. It is worth mentioning that we could detect ASIC current virtually in every neuron in the brain, suggesting a high distribution prevalence of ASICs in neurons. It is noteworthy that trigeminal nerves, where ASICs are expressed [42, 43], may also contribute to the acid-sensing of mice under acidic environments. Hence, the results that AC3 KO mice retained sniffing behaviors to the acidic Q-tip (Fig. 3) could be explained by two possible mechanisms: ASICs in OSNs stimulating the olfactory system, or activation of ASICs in the trigeminal nerve attracting the sniffing behavior.

Our results help explain previous findings and raise interesting considerations in the design of behavioral assessments. When neuroscientists conduct mouse behavioral studies, 5% acetic acid is commonly used between tests to neutralize odorants in the behavioral equipment to prevent the interfering effects of odor left by the previous animal. Inhaled table vinegar (with acetic acid concentrations varying from 4-8%) can temporarily numb our olfactory sensitivity for food smells. Here we also show that acetic acid interferes with normal mouse olfaction (Fig. 5). Why do mice exhibit reduced preference to attractive objects under an acidic volatile environment, and how do acidic volatiles interfere with normal olfaction? Our findings suggest that ASICs expressed in OSNs may play an important role. The mammalian olfactory system has anatomical logics for the perception of individual odorants and for sensory neural transmission into the brain [29, 44]. Each OSN expresses only 1 out of ∼1100 odor receptor genes [9, 45, 46] and the axons of individual OSNs project to two of 1800 glomeruli in the olfactory bulb [41, 47, 48], which further relay to the piriform cortex for information processing. Acids may interfere with regular odor perception through unselectively depolarizing different types of OSNs, which belong to different anatomical logic sets.

Although volatile acids may interfere with the perception of certain odorants, the actual effect may depend on the concentrations of acids. It is also possible that, on the contrary, slight depolarization of OSNs by mild activation of ASICs may promote olfactory sensitivity to sense regular odorants. Hence, table vinegar not only affects our taste bud, but also impacts our olfactory sensitivity for food smells. Our findings also help explain why vinegar has been used in some traditional medical practices. For instance, ancient Egypt and farmers in China used to spray vinegar around their homes in the hope of warding off bacteria or virus and preventing infectious diseases. One benefit of this practice was that volatile acetic acid could have intervened with the sensory communication of some pathogen-carrying animals including bats. Interestingly, this practice somewhat resembles the fact that environmental acidification disturbs ecological stability via interfering with animals’ olfactory communication and survival behaviors. Together, acidic volatiles can significantly impact animals’ olfaction and their behaviors, and one sensitive mediator for the acid-sensing is ASICs. ASIC activation may unselectively depolarize many subclasses of OSN, interfering with odor perception (Fig. 6). Considering humans and wildlife are more frequently exposed to acidic environments, more research is warranted to clearly elucidate the molecular and cellular mechanisms of acid-sensing in mammals and its ecological impacts it may have.

## Methods and Materials

### Mice

For EOG recording (Fig. 1&2 data), 3-chamber avoidance test, and Q-tip habituation/dishabituation test (Fig. 3 data), we used AC3 WT and KO mice with a C57BL/6J and 129 mixed background [12]. ASIC1 WT and KO MOE samples (Fig. 4 data) and mice used in 3-chamber preference test (Fig. 5 data) had C57Bl/6 background. Mice used in behavioral assays were group-housed and matched for age (8-18 weeks). Mice were singly housed for 5 days before tests. All mouse behavioral experiments were performed during the daytime light cycle. ASIC1 KO, AC3 KO and littermate control mice were bred from heterozygotes and genotyped as previously reported [12, 28, 49]. Mice were maintained on a 12 h light/dark cycle at 22°C and had access to food and water *ad libitum*. All animal procedures were approved by the Institutional Animal Care and Use Committee of the University of New Hampshire, the University of Washington, and the University of South Alabama, and performed in accordance with their guidelines.

### Electro-olfactogram (EOG) Recording

EOG recordings were performed as previously described with minor modifications [28, 50]. Briefly, after euthanization, the mouse head was bisected through the septum with a sharp razor blade and turbinates of MOE were exposed by removing the septal cartilage. Air puffs were applied to exposed MOE using a four-way slider air-puff valve controlled by an S48 Stimulator (Glass Technologies). Odorized or acidic volatile air was generated by blowing nitrogen through a horizontal glass cylinder that was half-filled either with TMT, odor mix, or volatile acids. The odor mix (dissolved in H_2_O) was comprised of eugenol, octanal, r-(+)-limonene, 1-heptanol, s-(−)-limonene, acetophenone, carvone, 3-heptanone, 2-heptanone, ethyl vanillin, and citralva, each at 50 μM. The air puff duration was 200 milliseconds. The tip of the puff application tube was directly pointed to the recording site. The flow rate was 1 L/min and EOG recording sites were in the turbinate II of the MOE [28, 50].

A filter paper immersed in Ringer’s solution (in mM, 125 NaCl, 2.5 KCl, 1 MgCl_2_, 2.5 CaCl_2_, 1.25 NaH_2_PO_4_, 20 HEPES, and 15 D–glucose, with pH 7.3 and osmolarity 305 mOsm/L) was used to hold the sample on a plastic pad during recording [28, 50]. The filter paper was connected to the recording circuit, as the ground electrode was immersed in Ringer’s bath solution. Electrophysiological field potential was amplified with a CyberAmp 320 (Molecular Devices) and digitized by a Digidata 1332A processor at 10 kHz as well as simultaneously by a MiniDigi 1A processor at 1 kHz (Molecular Devices) [28, 50]. The signals were acquired on-line using software pClamp 10.3 (Molecular Devices) in combination with Axoscope 10 (Molecular Devices). Forskolin/IBMX and diminazene were applied in Ringer’s solution to the surface of MOE, respectively. These chemicals were washed away using Ringer’s solution (twice). Since residual liquid on the MOE surface prevented EOG recording, a layer of filter paper was put onto the nasal cavity to drain liquid away. Afterwards, MOE responded to air puff stimulation again and EOG signals re-appeared.

### Q-tip habituation/dishabituation test in home cage

All tests were done in home cages, where the testing mouse has been singly housed for 5 days. Odor stimulations were delivered with a cotton-tipped swab placed through the cage top ∼8 cm above the bedding. After 10 min of habituation with a cotton-tipped swab without odor stimulant, the test mouse was stimulated by several applications: water, TMT odor, and 5% acetic acid. Each stimulus was 2 min in duration with 1 min inter-trial interval. The sequence of the odor stimulation was as follows: water1, water2, water3, water4, TMT odor5, water6, and water1, water2, water3, water4, 5% acetic acid5, water6. Time spent sniffing the Q-tip was measured by manual observation with a stopwatch. Sniffing was only scored when the test mouse’s nose was close from and pointing to the swab. Biting of the swab by the mouse was excluded.

### Immunofluorescence staining

Mice were deeply anesthetized by intraperitoneal (IP) injection of ketamine (100 mg/kg), and then perfused transcardially with ice-cold 0.1 M phosphate buffer saline (PBS) followed by 4% paraformaldehyde in 0.1 M PBS. The nasal bone, including all olfactory tissues, was incubated overnight at 4 °C in 4% paraformaldehyde and then decalcified by exposure to 500 mM EDTA in 0.1 M PBS (pH 7.5) for 48 h at 4 °C, after which the MOE was washed by 0.1 M PBS for 10 minutes for 3 times, and then dehydrated by 30% sucrose in 0.1 M PBS for 24 h at 4°C, and finally embedded in O.C.T resin before being sectioned at −18 ° C using a cryostat to a thickness of 30 μm according to standard procedures. MOE sections mounted on gelatin coated slices were washed three time with PBST, blocked and then first incubated with primary antibody overnight at 4 ° C in blocking buffer (PBS containing 10% normal goat serum (vol/vol), 2% bovine serum albumin (weight/vol), 0.2% TritonTM X-100 (vol/vol)), washed three times in PBST (PBS with 0.2% Triton X-100, vol/vol), and then incubated with secondary antibodies for one hour at room temperature. Primary antibodies included mouse anti-ASIC1 (1:300, UC Davis, #75-277), rabbit anti-AC3 (1:500, Santa Cruz biotechnology Inc, #SC588) or rabbit anti-AC3 (1:20000, EnCor biotechnology Inc, #RPCA-ACIII), mouse anti-ARL13B (1:300, NeuroMab, #75-287). Secondary antibodies were Alexa fluor 488-, 546- or 648-conjugated (Invitrogen, La Jolla, CA). Finally, sections were counterstained with DAPI (SouthernBiotech™, #OB010020) and images were acquired with confocal microscope (Nikon, A1R-HD). To estimate the distribution percentage of ASIC1-positive OSNs in the MOE, we used Fiji-ImageJ software to count ASIC1-positive neurons (merged with AC3 signals in the cell body) from 9 randomly chosen areas (out of 3 mice) in the MOE to obtain an average of ASIC1-positive neurons per mm^2^, and then divided by 97,000, which is an average density of mature receptor cells per mm^2^ in mouse MOE surface area [51].

### 3-chamber acid-interference olfactory test (and 3-chamber TMT avoidance test)

The apparatus for acid-interference olfactory test was a rectangular, three-chamber box. Each chamber was 22 × 20 × 13 cm and the walls of chamber were made from Plexiglas. There were open holes between chambers that allowed a subject mouse to freely explore each chamber. For subject male mice, identical small pieces of clean cotton nestlet moisturized with 20 µl of double-distilled water were first placed in the right and left chambers respectively, and then the mouse was placed in the middle chamber and let to habituate in the apparatus for 5 minutes. After the habituation, one cotton nestlet was replaced by an identical size of the cotton nestlet which came from a cage housing female mice. Then both cotton nestlets were moistened with water drops (20 µl). The exploration lasted 10 minutes and was video recorded. We further tested the subject mice in several different environments. For each test, all procedures were the same except that water drops were replaced by 5mM ethyl vanillin, 5% acetic acid, and 10% acetic acid, during both habituation and exploration. For testing the subject female mice in different environments, all procedures were the same except peanut butter was used as an attractive object to replace the cotton with female odorants for male mice. We used water and 5% acetic acid, respectively, for each test. The exploration time in each chamber was video-typed and analyzed offline with EthoVision XT software (Noldus). The 3-chamber TMT avoidance test was conducted in a similar manner as the 3-chamaber acid-interference olfactory test, with odor changed to TMT.

### Data analysis and graphic data presentation

Data were analyzed with Clampfit 10.3, Microsoft Excel and GraphPad Prism. The decay phases of acid-evoked EOG field potential were fitted with a mono-exponential function *f*(*t*) = *A*_0_* exp(−*t*/τ) + *a*, where τ is the decay time constant, *A*_0_ is the maximal response, and *a* is the residual response. When appropriate, statistical analyses were conducted using ANOVA for multiple group comparison or Student’s t-test with a two-tailed distribution. n.s. not significant, * *p* < .05, ** *p*< .01, *** *p* < .001. Data was considered as statistically significant if *p* < .05 and values in the graph are expressed as mean ± standard error of the mean.

## Acknowledgements

We are grateful to Dr. Daniel Storm at the University of Washington, who has provided research support and resources for completing part of the experiments. We thank the members of the Chen Laboratory for their critical review of the manuscript and Dr. Tao Wang for preparing ASIC1 wildtype and knockout samples. This study was supported by National Institutes of Health Grants MH105746, AG054729 and GM113131 to X.C.; Cole Neuroscience and Behavioral Faculty Research Awards to X.C.; and UNH Summer TA Research Fellowships to J.Y and M.S.

